# Structure of the large extracellular loop of FtsX and its interaction with the essential peptidoglycan hydrolase PcsB in *Streptococcus pneumoniae*

**DOI:** 10.1101/432344

**Authors:** Britta E. Rued, Martín Alcorlo, Katherine A. Edmonds, Siseth Martínez-Caballero, Daniel Straume, Yue Fu, Kevin E. Bruce, Hongwei Wu, Leiv Sigve Håvarstein, Juan A. Hermoso, Malcolm E. Winkler, David P. Giedroc

## Abstract

*Streptococcus pneumoniae* is a leading killer of infants and immunocompromised adults and has become increasingly resistant to major antibiotics. Therefore, the development of new antibiotic strategies is desperately needed. Targeting bacterial cell division is one such strategy, specifically targeting essential proteins for the synthesis and breakdown of peptidoglycan. One complex important to this process is FtsEX. FtsEX comprises an integral membrane protein (FtsX) and cytoplasmic ATPase (FtsE) that resembles an ATP-binding cassette (ABC) transporter. Here, we present NMR solution structural and crystallographic models of the large extracellular domain of FtsX, denoted ECL1. The structure of ECL1 reveals an upper extended β-hairpin and a lower α-helical lobe, each extending from a mixed α-β core. The helical lobe mediates a physical interaction with the peptidoglycan hydrolase PcsB, via the coiled-coil domain of PcsB (PcsB-CC). Characterization of *S. pneumoniae* D39 derived strains harboring mutations in the α-helical lobe shows that this subdomain is essential for cell viability and required for proper cell division of *S. pneumoniae*.

**IMPORTANCE:** FtsX is a ubiquitous bacterial integral membrane protein involved in cell division that regulates the activity of peptidoglycan (PG) hydrolases. FtsX is representative of a large group of ABC3 superfamily proteins that function as “mechanotransmitters,” proteins that relay signals from inside to the outside of the cell. Here we present a structural characterization of the large extracellular loop (ECL1) of FtsX from the human opportunistic pathogen *Streptococcus pneumoniae*. We show a direct interaction between the peptidoglycan hydrolase PcsB and FtsX, and demonstrate that this interaction is essential for cell viability. As such, FtsX represents an attractive, conserved target for the development of new classes of antibiotics.

## INTRODUCTION

*Streptococcus pneumoniae* is a Gram-positive, opportunistic respiratory pathogen (1–3) that has acquired antibiotic resistance worldwide (4–6). This ovococcal bacterium relies on highly conserved cell wall machinery to divide and grow (7, 8). The cell wall is primarily composed of peptidoglycan (PG), a macromolecule composed of repeating subunits of N-acetylglucosamine and N-acetylmuramic acid linked by PG peptide side chains (9, 10). Regulation of the synthesis and remodeling of PG is essential for bacterial growth and viability, due to the turgor pressure bacterial cells must withstand (10–12). One vital process for the synthesis of PG is the controlled insertion of new strands of PG. This process requires timed cleavage of the old PG matrix to allow incorporation of new nascent strands (13). PG hydrolases are the primary enzymes that carry out PG cleavage and remodeling (14, 15). Thus, regulation of these hydrolases and activation at specific times during the cell cycle is required for proper cell growth. Specific protein complexes are utilized by bacterial cells to regulate these enzymes. This work focuses on understanding the structure and function of one of these protein complexes.

From *Mycobacterium tuberculosis* to *Caulobacter crescentus*, the ABC-transporter-like protein complex FtsEX acts as a key regulator of PG hydrolysis and divisome assembly (16–19). The proposed mechanism of FtsEX activation of PG hydrolases is as follows. FtsE, upon sensing an unknown signal from inside the cell, hydrolyzes ATP to ADP. Hydrolysis causes a conformational change transmitted through the membrane via FtsX, an integral membrane protein with two extracellular loops, denoted the large (ECL1) and small (ECL2) loops. These extracellular loops interact with either the cell wall hydrolases or with effector proteins, which results in activation of PG hydrolysis via an unknown mechanism (16, 18, 20–25). In *E. coli*, it has been demonstrated that FtsX interacts with the effector protein EnvC to activate the PG amidases AmiA and AmiB (24, 25). In addition, FtsX interacts with other division proteins such as FtsA, where it regulates the polymerization of FtsA and recruitment of downstream division proteins (26). In other organisms including *B. subtilis* and *M. tuberculosis*, FtsEX also activates PG hydrolases (16, 23). Interestingly, FtsEX is non-essential in rod-shaped bacteria such as *E. coli* and *B. subtilis* (23, 24, 26–28). However, in *S. pneumoniae* FtsEX is absolutely essential (21) and depletion of FtsEX results in cell rounding and cessation of growth (20, 21).

In the case of *S. pneumoniae*, genetic experiments suggest that both outward-facing domains of FtsX, ECL1 and ECL2, interact with the essential PG hydrolase PcsB, via its long coiled-coil (CC) domain (20, 21). However, there is little direct biochemical evidence for this interaction. ECL1 and ECL2 are thought to allosterically activate the catalytic activity of cysteine, histidine-dependent amidohydrolase/peptidase (CHAP) domain of PcsB (20). The crystal structure of full-length PcsB including the CC domain, an alanine-rich linker region, and the CHAP domain provides insight into the mechanism of how this may occur (22). While the PcsB structure implies FtsEX activates PcsB by displacing the catalytic domain from the CC domain, the exact nature of the FtsX::PcsB interaction remains unknown.

In order to understand how FtsX activates PcsB, we determined the structure of the large extracellular loop of FtsX (FtsX_ECL1_) by both multidimensional NMR spectroscopy and X-ray crystallography. FtsX_ECL1_ harbors a conserved mixed α-β core and a lower α-helical lobe extending from the core identified previously in *M. tuberculosis* FtsX (16), and an extended β-hairpin that is unique to *S. pneumoniae* FtsX_ECL1_. The N-terminal β1 and C-terminal β6 strands are adjacent in the core and connect ECL1 to the TM1 and TM2 helices, respectively, in the membrane. PcsB-CC-mediated chemical shift perturbations of ^1^H-^15^N HSQC spectra of FtsX_ECL1_ reveal that the helical lobe consisting of the α2 helix and the α2-β5 linker (residues 107–134) of FtsX_ECL1_ interacts with PcsB-CC. To determine if this interaction is required for FtsX function in bacterial cells, we constructed a merodiploid strain that allows for conditional expression of the mutant *ftsX*. We demonstrate that specific amino acid substitutions in the FtsX-PcsB interface are lethal or cause pronounced morphological defects despite the fact these FtsX_ECL1_ mutant proteins are expressed at near wild-type levels. These findings support the model that a direct physical interaction between FtsX and PcsB is required for activation of PcsB PG hydrolytic activity.

## RESULTS

### The three-dimensional structure of FtsX_ECL1_

The three-dimensional structure of FtsX_ECL1_ (residues 46–168) was solved by both NMR spectroscopy (Fig. 1A) and X-ray crystallography (Fig. 1B). The folded structure (residues 57–166) reveals a central core composed of a four-stranded antiparallel β-sheet (β1, β6, β4, β5) and two helices (α1 and α3), an α-helical lobe (residues 107–135) harboring the α2 helix, and an extended β-hairpin (β2, β3). The β-hairpin and helical lobes are connected to the central core by hinges. Details for structural determination of FtsX_ECL1_ by NMR are presented in the methods section and structure statistics are summarized in Table 1. The solution structure shows that while the central mixed α-β core adopts a well-defined conformation, the two appended lobes are highly dynamic on multiple timescales (*vide infra*), presenting a range of conformations among the 20 members of the FtsX_ECL1_ NMR structural ensemble (Fig. 1A).

**TABLE 1.** Structural statistics for NMR solution structure of FtsX_ECL1_ (From the ensemble of 20 best NMR structures).

|  |  |
| --- | --- |
| **NMR structure (PDB code)** | 6MK7 |
| **(BMRB code)** | 30523 |
| **NMR distance and angle restraints** |  |
| Total NOE-based distance constraints | 1711 |
| intraresidue (i=j) | 232 |
| sequential ($\vert i-j\vert = 1$) | 499 |
| medium-range ($1 < \vert i-j\vert < 5$) | 365 |
| long-Range ($\vert i-j\vert \geq 4$) | 615 |
| Maximum distance violation | 0.49 Å |
| Dihedral angle restraints | 110 |
| Maximum dihedral angle violation | 7° |
| Total number of RDCs used (measured) | 82 (112) |
| Q factor | 0.11 |
|  | (0.39) |
| Correlation (experimental to calculated) | 0.99 |
|  | (0.89) |
| **RMS deviations from idealized covalent geometry** |  |
| Covalent Bond Lengths (Å) | 0.011 Å |
| Covalent Angle Values (°) | 1.3 ° |
| **Ramachandran analysis <sup>a,b</sup>** |  |
| Most favored (%) | 86.5 |
| Additionally allowed (%) | 8.4 |
| Generously allowed (%) | 1.8 |
| Disallowed (%) | 3.3 |
| **RMSD Values** |  |
| Backbone atoms (all) | 1.5 Å |
| All heavy atoms (all) | 1.9 Å |
| Backbone atoms (core<sup>a</sup>) | 0.4 Å |
| All heavy atoms (core<sup>a</sup>) | 0.9 Å |

<sup>a</sup>Calculated over the core of the structure, excluding the helical lobe and β-hairpin, residues 59-65, 89-108, 134-165. <sup>b</sup>As computed by Procheck.

**Figure 1:**
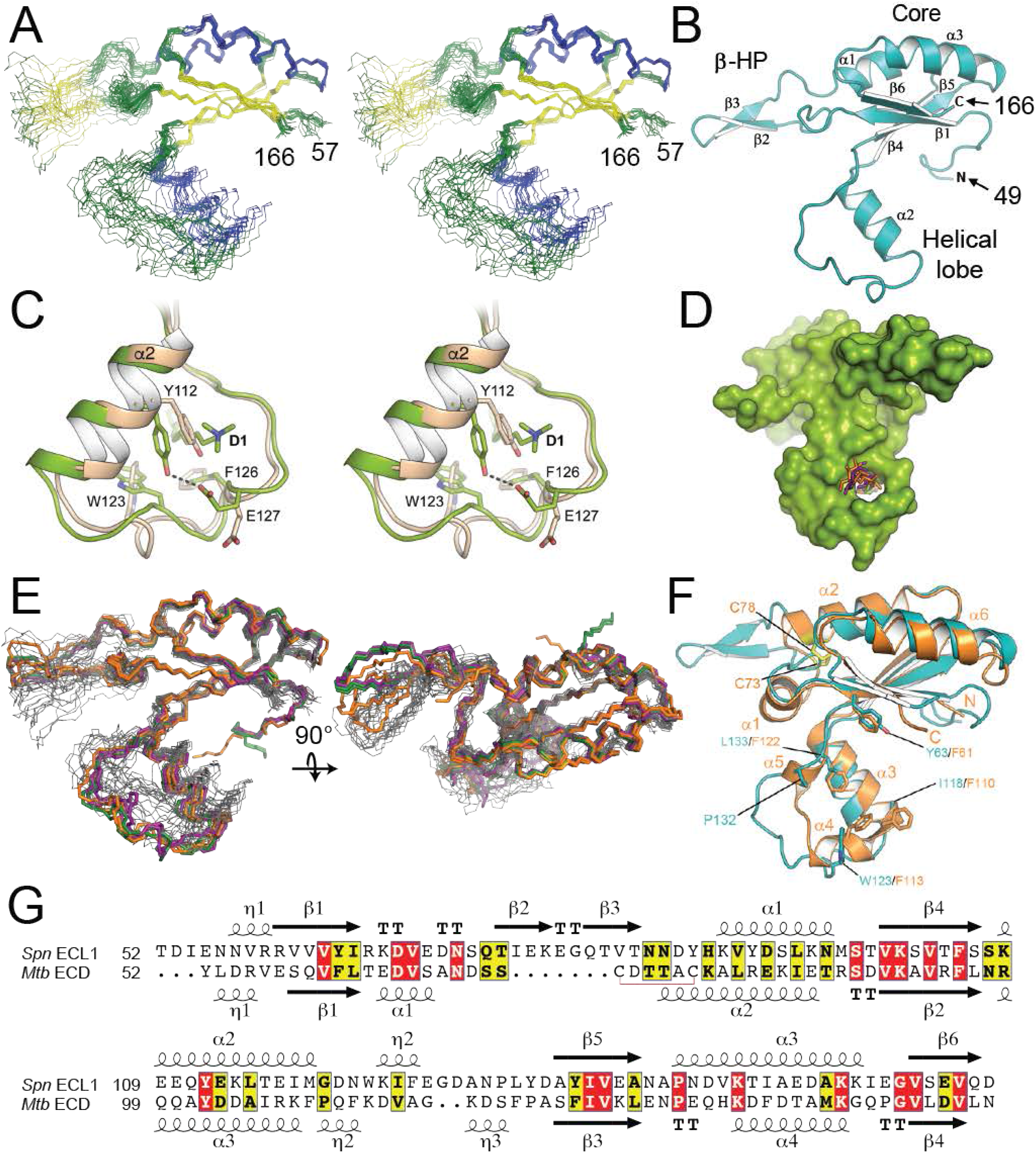
The structure of FtsX_ECL1_ from *Streptococcus pneumoniae.* A) Stereo view of the 20 conformers of the FtsX_ECL1_ NMR structure as backbone traces, with helices shown in blue and (3-strands shown in yellow. N- and C-termini of the domain are indicated by the residue numbers 57 and 166, respectively. B) Cartoon representation of the FtsX_ECL1_ structure obtained by X-ray crystallography (chain B in FtsX_ECD_-D1) in which the different secondary structure elements have been numbered and labeled. N- and C-termini of the domain are indicated and correspond to residue numbers 49 and 166, respectively. C) Stereo view showing changes in helical lobe upon interaction with detergent 1. Apo-form (chain B) is colored in pale brown and holo-form (chain A) is colored in green. Relevant residues affected by the presence of the detergent are depicted in capped sticks. Polar interaction represented by dashed line. D) Surface representation of the FtsX_ECL1_ crystal structure in which the three different detergent molecules are superimposed and shown in sticks (detergent 1, 2 and 3 are colored in green orange and purple, respectively). E) An overlay of the backbone traces for the six FtsX_ECL1_ structures (in black) with the backbone traces of the 20 FtsX_ECL1_ NMR conformers (in green) F) Structural superposition of the crystal structure of FtsX_ECL1_ from S. *pneumoniae* obtained by crystallography (colored in cyan) and the crystal structure of FtsX_ECD_ from *M. tuberculosis* (PDB 4N8N, colored in orange). Secondary structure elements observed in the *M. tuberculosis* structure are labeled. Some residues in both structures are shown as sticks and numbered (see main text). Cys residues involved in disulfide bond formation in *M. tuberculosis* FtsX_ECD_ are colored in yellow. See main text for details. G) Sequence alignment of ECL1 domains from S. *pneumoniae* (Spn ECL1) and *M. tuberculosis* (Mtb ECD). Secondary structure elements are indicated and numbered.

Three different structures, with resolutions ranging from 2.0 to 2.3 Å, were solved by X-ray crystallography, each in the presence of a different detergent. The presence of detergents was critical as in their absence crystals diffracted at very low resolution (≤4 Å), suggesting significant mobility in some protein regions. Details for the crystallographic determination are provided in the methods section and structure statistics summarized in Table 2. In all cases, two protein molecules are present in the asymmetric unit (Fig. S1), yielding a total of six independent structures and sufficient electron density for both the protein structure and associated detergent molecules (Fig. S2). Different conformations were observed for the β-hairpin and the helical lobe among the six structures depending on the presence and identity of the bound detergent molecule to FtsX_ECL1_ (Fig. S1B). The structural variations observed in these crystallographic structures, however, are less dramatic than those observed in the NMR conformer bundle, obtained in the absence of ligand, although the regions with high variability are similar.

**TABLE 2.**
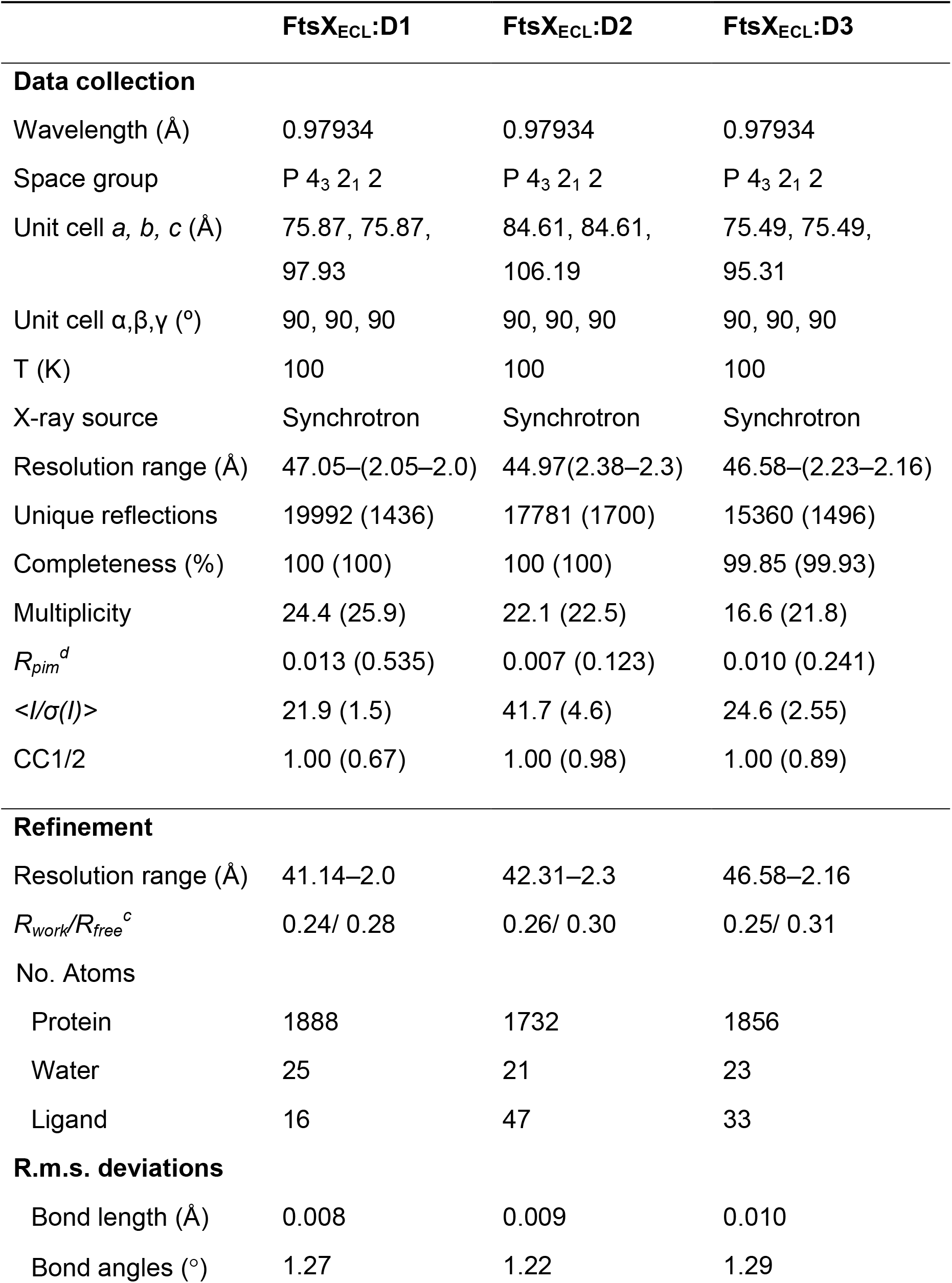
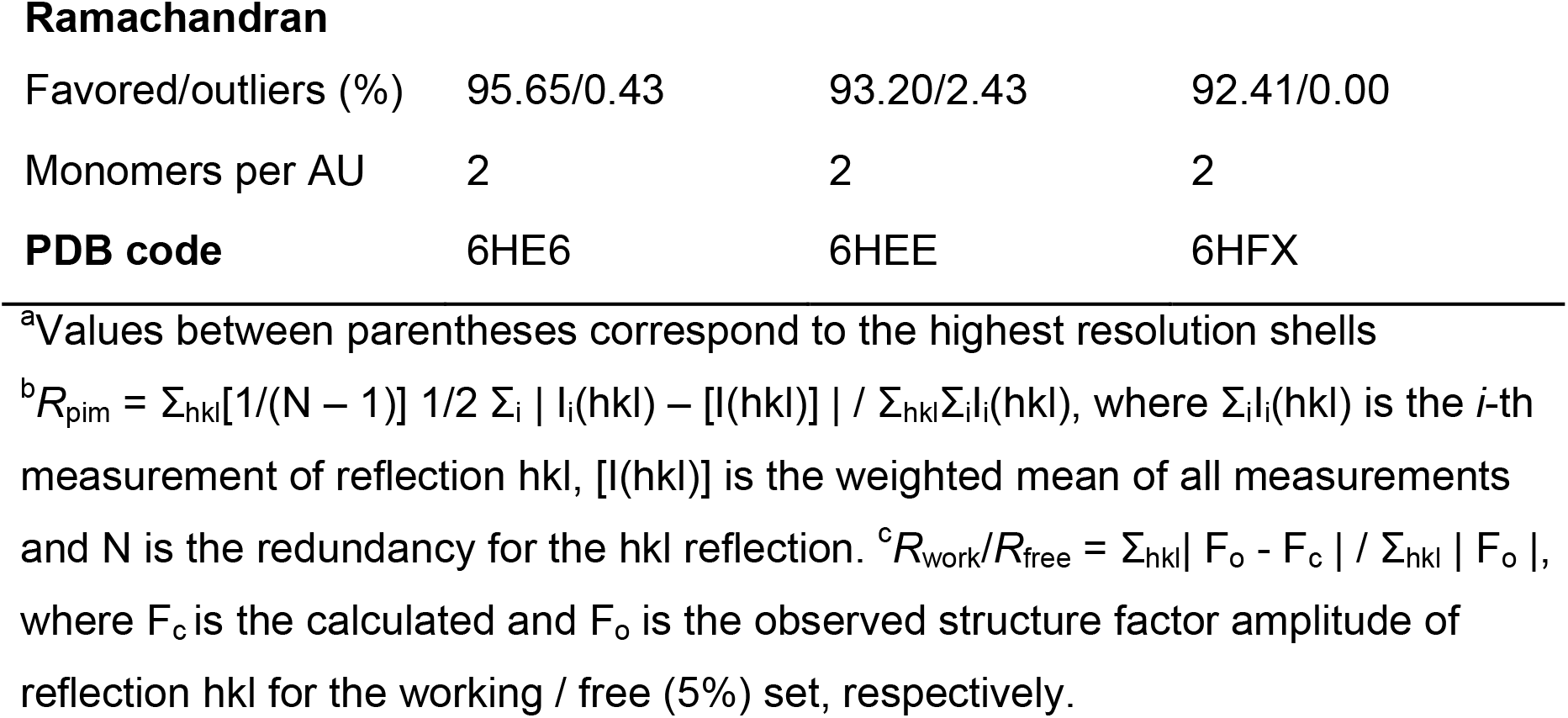
Crystallographic data collection and refinement statistics.^a^

|  | **FtsX<sub>ECL</sub>:D1** | **FtsX<sub>ECL</sub>:D2** | **FtsX<sub>ECL</sub>:D3** |
| --- | --- | --- | --- |
| **Data collection** |  |  |  |
| Wavelength (Å) | 0.97934 | 0.97934 | 0.97934 |
| Space group | P 4<sub>3</sub> 2<sub>1</sub> 2 | P 4<sub>3</sub> 2<sub>1</sub> 2 | P 4<sub>3</sub> 2<sub>1</sub> 2 |
| Unit cell *a*, *b*, *c* (Å) | 75.87, 75.87,<br>97.93 | 84.61, 84.61,<br>106.19 | 75.49, 75.49,<br>95.31 |
| Unit cell $\alpha, \beta, \gamma$ (°) | 90, 90, 90 | 90, 90, 90 | 90, 90, 90 |
| T (K) | 100 | 100 | 100 |
| X-ray source | Synchrotron | Synchrotron | Synchrotron |
| Resolution range (Å) | 47.05–(2.05–2.0) | 44.97(2.38–2.3) | 46.58–(2.23–2.16) |
| Unique reflections | 19992 (1436) | 17781 (1700) | 15360 (1496) |
| Completeness (%) | 100 (100) | 100 (100) | 99.85 (99.93) |
| Multiplicity | 24.4 (25.9) | 22.1 (22.5) | 16.6 (21.8) |
| $R_{pim}^d$ | 0.013 (0.535) | 0.007 (0.123) | 0.010 (0.241) |
| $\langle I/\sigma(I) \rangle$ | 21.9 (1.5) | 41.7 (4.6) | 24.6 (2.55) |
| CC1/2 | 1.00 (0.67) | 1.00 (0.98) | 1.00 (0.89) |
| **Refinement** |  |  |  |
| Resolution range (Å) | 41.14–2.0 | 42.31–2.3 | 46.58–2.16 |
| $R_{work}/R_{free}^c$ | 0.24/ 0.28 | 0.26/ 0.30 | 0.25/ 0.31 |
| No. Atoms |  |  |  |
| Protein | 1888 | 1732 | 1856 |
| Water | 25 | 21 | 23 |
| Ligand | 16 | 47 | 33 |
| **R.m.s. deviations** |  |  |  |
| Bond length (Å) | 0.008 | 0.009 | 0.010 |
| Bond angles (°) | 1.27 | 1.22 | 1.29 |
| Ramachandran |  |  |  |
| Favored/outliers (%) | 95.65/0.43 | 93.20/2.43 | 92.41/0.00 |
| Monomers per AU | 2 | 2 | 2 |
| **PDB code** | 6HE6 | 6HEE | 6HFX |

<sup>a</sup>Values between parentheses correspond to the highest resolution shells
<sup>b</sup> $R_{\text{pim}} = \sum_{\text{hkl}} [1/(N - 1)]^{1/2} \sum_i |I_i(\text{hkl}) - [I(\text{hkl})]| / \sum_{\text{hkl}} \sum_i I_i(\text{hkl})$ , where $\sum_i I_i(\text{hkl})$ is the $i$ -th measurement of reflection hkl, $[I(\text{hkl})]$ is the weighted mean of all measurements and N is the redundancy for the hkl reflection. <sup>c</sup> $R_{\text{work}}/R_{\text{free}} = \sum_{\text{hkl}} |F_o - F_c| / \sum_{\text{hkl}} |F_o|$ , where $F_c$ is the calculated and $F_o$ is the observed structure factor amplitude of reflection hkl for the working / free (5%) set, respectively.

In any case, the crystal structures suggest that changes in the protein backbone and side chains of the helical lobe occur when a detergent ligand is bound (Fig. 1C). These changes create a cavity in which the detergent molecules insert (Fig. 1D), with a large part of the helical lobe affected by the interaction with detergent (from Q111 to E127) (Fig. 1C). Residues Y112, W123, E127 and F126 are strongly perturbed upon detergent ligand binding (Fig. 1C) with the hydroxyl moiety of Y112 interacting with carboxylate of E127. Changes in W123 and F126 stabilize the hydrophobic region of detergent 1. A similar interaction pattern is observed for detergents 2 and 3; additional interactions are observed with N131 with detergents 2 and 3, which are characterized by a larger hydrophilic head (maltose) (Fig. S1E-F). Although a physiological role for detergent binding to the helical lobe is unknown, many of these same residues are important for the interaction with PcsB (*vide infra*).

A structural comparison with *M. tuberculosis* FtsX_ECL1_ (16) (PDB 4N8N) reveals differences in both the overall structure (RMSD value of 2.2 Å) and in the appended lobes of the core domain (Fig. 1F). The main differences between the mycobacterial and pneumococcal FtsX_ECL1_ domains are the presence of an extra helix (α1) and a disulfide bond in *M. tuberculosis* ECL1 that are absent in pneumococcal ECL1, and an extended β-hairpin (residues 71–87) unique to the pneumococcal ECL1 domain. It is worth noting that in *M. tuberculosis* ECL1, the upper and lower lobes form a large hydrophobic cleft with four exposed Phe residues (F61, F110, F113 and F122) and this region was suggested as a strong candidate interaction surface between FtsX and RipC (16). These phenylalanine residues are not conserved in the pneumococcal ECL1 (Fig. 1F-G) but the hydrophobic nature of this region is preserved (Y63, I118, W123 and L133).

### FtsX_ECL1_ is dynamic in solution

We next performed additional NMR experiments to explore the mobility of the FtsX_ECL1_ domain, both to validate the heterogeneity of the structural ensemble in solution, and to elucidate function. The ^1^*D*_HN_ residual dipolar couplings (RDCs) obtained by weak alignment in Pf1 filamentous phage correspond well to previously determined secondary structure elements (29), with uniform values for the entire length of α1 and α2, as expected for straight helices. In contrast, ^1^*D*_HN_ values near 0 for the N-terminal tail, the very C-terminus, and the non-helical part of the helical lobe between α2 and β5, are suggestive of significant conformational disorder in solution (Fig. S3A). These regions of small or zero RDCs are regions of very high RMSD in the NMR structure bundle (Fig. S3B). As anticipated, the correlation between experimentally measured RDCs and predicted RDCs back-calculated from the structures (30) is high, but only in the core subdomain, and match poorly in the β-hairpin for most of the crystal structures (Fig. S3C-D).

We previously reported that the ^15^N-{^1^H} heteronuclear NOE (hNOE) is low or negative at the termini, indicating that they are highly flexible in solution (29). The hNOE is also smaller in the β2-β3 hairpin region, as well as in the C-terminal end of the α2 helix and the subsequent coiled region leading into β5 (29) (Fig. S3E). Mapping these dynamics data onto a representative structure from the solution NMR ensemble shows that these regions with fast-timescale dynamics correspond to the β-hairpin and the helical lobe (Fig. S3F). Information on ps-ns fast-timescale motions extracted from the *R*_2_/*R*_1_ ratio also reveals that α2-β5 linker is highly flexible, while the β1-β2 linker and β-hairpin show an elevated *R*_2_/*R*_1_ ratio, specifically indicative of millisecond-timescale slow conformational exchange (Fig. S3G-H). These findings were directly confirmed by (Carr-Purcell-Meiboom-Gill) relaxation dispersion NMR spectroscopy (31) (Fig. S3I-J). We conclude that the β-hairpin exhibits flexibility on both the sub-ns and ms timescales. Interestingly, the β1-β2 linker also shows increased B-factors that are qualitatively consistent with ms-timescale conformational exchange observed by NMR.

These complex motions observed in the solution dynamics experiments are reflected in the heterogeneity of the NMR structure, with high Cα RMSDs particularly in the α2-β5 linker but also in the β-hairpin and β1-β2 linker, thus validating the conformational spread in the ECL1 structure in solution (Fig. S3B). The dynamic nature of the helical lobe is also reflected in the heterogeneity and B-factors of the crystal structures (Fig. S3K-L). Full-length FtsX itself is likely a dimer *in vivo* (32), and one can speculate that the flexible helical lobe and β-hairpin regions may contribute to dimerization or to interactions with a binding partner such as PcsB.

### The PcsB coiled-coil (PcsB-CC) domain interacts with FtsX_ECL1_

The ^1^H-^15^N HSQC spectrum for ECL1 has excellent chemical shift dispersion and lends itself readily to studies of protein-protein interactions (Fig. S4A). In contrast, full-length PcsB is 42 kDa and forms a dimer, and is thus challenging to study by NMR due to its size. We therefore constructed truncation mutants of PcsB focusing on the coiled-coil domain (PcsB-CC), thus limiting the molecular weight to 23–24 kDa. ^15^N labeled PcsB-CC (47–267) and PcsB-CC (47–254) are both characterized by ^1^H-^15^N HSQC spectra with limited ^1^H signal dispersion (Fig. S6A), consistent with the high helical content of this domain. Circular dichroism confirms that these proteins are primarily α-helical, in agreement with the crystal structure (22), thus indicating that they are properly folded (Fig. S6A) and can be used for ECL1 binding studies.

To determine if PcsB physically interacts directly with FtsX_ECL1_, we titrated unlabeled PcsB-CC (47–267) into ^15^N labeled FtsX_ECL1_ at molar ratios of 1:1, 2:1, 4:1, and 6:1, respectively, and recorded ^1^H-^15^N HSQC spectra (Fig. S4A-B). Numerous crosspeaks move in response to the addition of PcsB-CC to ^15^N FtsX_ECL1_. The largest changes occur in the helical lobe of FtsX_ECL1_ (Fig. 2A, S4-S5). In particular, residues M119, W123, I125, F126, G128 exhibit the greatest changes in crosspeak position and intensity in the 1:2 FtsX_ECL1_ to PcsB-CC sample (Fig. 2A, Fig. S4C); when additional PcsB-CC is added, these crosspeaks broaden beyond detection (Fig. S5B-C). A reciprocal ^1^H-^15^N HSQC experiment with ^15^N labeled PcsB-CC (47–254) further confirms an interaction with unlabeled FtsX_ECL1_ (Fig. S6B), as multiple crosspeaks shift upon the addition of increasing FtsX_ECL1_ (Fig. S6B). We measured the binding affinity of the FtsX_ECL1_-PcsB-CC complex using isothermal titration calorimetry (ITC), which reveals a *K*_a_ of 3.0 × 10^4^ M^−1^ (*K*_d_ ~ 34 µM) (Table 3, Fig. 2B). These data provide the first biochemical evidence for a direct physical interaction between PcsB-CC and FtsX_ECL1_ (Fig. 2, Fig. S4-S6).

**TABLE 3.** Thermodynamic parameters of wild-type and mutant FtsX_ECL1_ from direct analysis of isothermal titration calorimetry (ITC).^a^

| Experiment | Protein | $K_A$ ( x 10<sup>4</sup> M<sup>-1</sup>) | $K_D$ (μM) | $\Delta H$ (kcal<br>mol<sup>-1</sup>) | $\Delta S$ (kcal<br>mol<sup>-1</sup> K<sup>-1</sup>) |
| --- | --- | --- | --- | --- | --- |
| ITC | FtsX<sub>ECL1</sub> | 3.0 ± 0.5 | 34 ± 6 | -2.1 ± 0.2 | 13.4 |
| ITC | L115A, M119A<br>FtsX<sub>ECL1</sub> | N/A<sup>b</sup> | N/A<sup>b</sup> | N/A<sup>c</sup> | N/A<sup>c</sup> |

<sup>a</sup> Solution conditions: 50 mM sodium phosphate, 50 mM NaCl, 0.5 mM EDTA, pH 7.0 at 25 °C. The chemical model used to fit the data is indicated in *Supplemental Materials* *and Methods*. <sup>b</sup>No observed heat.

**Figure 2:**
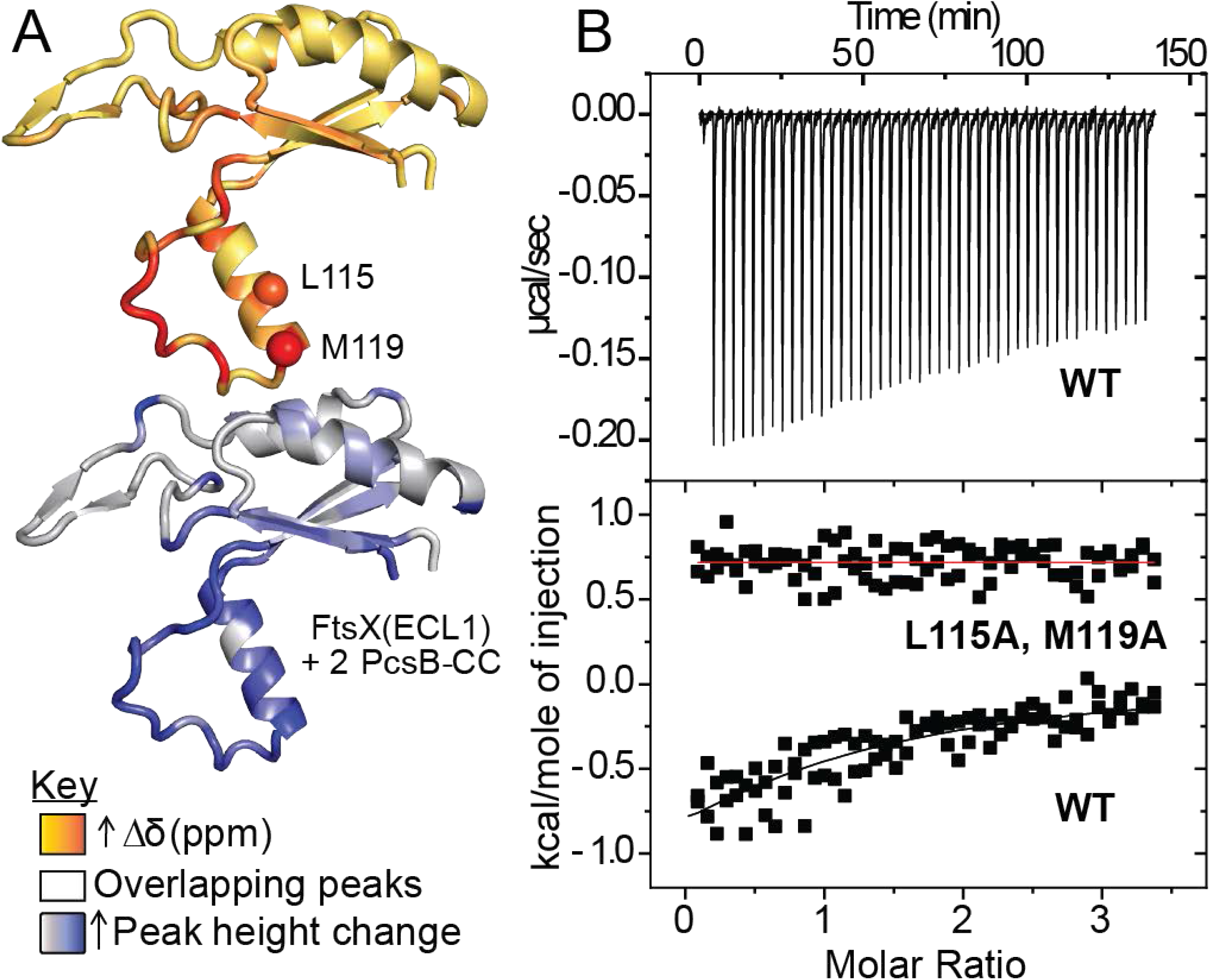
FtsX_ECL1_ binds PcsB-CC. A) Significant chemical shifts and peak height changes upon ^1^H-^15^N HSQC titration of 50 μM ^15^N FtsX_ECL1_ with 100 μM unlabeled PcsB-CC map to the lower lobe of FtsX_ECL1_. Chemical shift (Δδ, ppm) and peak height changes are mapped as color gradients on the FtsX_ECL1_ structure, orange to red and light grey to blue, respectively. L115 and M119 a carbons are shown as spheres on the upper image. Peaks that overlap in the^1^H-^15^N HSQC spectra are colored white. Chemical shift and peak height change upon addition of 2 molar equivalents of PcsB-CC to FtsX_ECL1_ are mapped to the structure. B) Representative titration of PcsB-CC with wild-type (WT) FtsX_ECL1_ or L115A/M119A FtsX_ECL1_ as monitored by ITC. Conditions: 50 mM potassium phosphate, 50 mM NaCl, 0.5 mM EDTA, pH 7.0 at 25.0 °C. Top panel, corrected ITC data; bottom panel, kcal/mole of injection vs. molar ratio. The black line overlapping the WT data indicates the best fit to a one-site binding model. Fitting parameters are summarized in Table 3. A red line drawn through the L115A/M119A FtsX_ECL1_ data is for reference.

### The interaction region of FtsX_ECL1_ with PcsB-CC is essential for cell growth and proper morphology

Having identified the interaction region between FtsX_ECL1_ and PcsB-CC, we next sought to determine the degree to which this interaction interface mapped by NMR spectroscopy contributes to pneumococcal viability. A multiple sequence alignment of this region (residues 102–155) among bacterial species in which FtsX has been studied and in related Streptococcal species (Fig. S7A) reveals that amino acids in this region are either partially or completely conserved (Fig. S7A). We therefore decided to target E109, Q111, L115, M119, W123, F126, and N131 singly or in combination for substitution with alanine (Fig. S7A). Given the essentiality of the FtsX_ECL1_ and PcsB-CC interaction, we predicted that mutating these residues might be lethal (20). To allow for the cross-in of potentially lethal point mutations, we employed the Janus cassette method to insert point mutations at the native site of *ftsX* into a strain containing an ectopic copy of *ftsX*^+^under a zinc inducible promoter (33) (Fig. S7B). We then transformed markerless mutant alleles of *ftsX* in the presence of zinc. This allows for expression of the wild-type *ftsX*^+^and mutant *ftsX* simultaneously. As long as the mutant *ftsX* is not dominant negative, we could obtain a strain that expresses the wild-type copy of *ftsX*^+^under zinc induction and mutant *ftsX* only in the absence of zinc (Fig. S7B).

Zinc toxicity has been observed to cause aberrant cell morphology and growth inhibition in *S. pneumoniae*, when cells are not supplemented with manganese (34, 35). To rule out any deleterious effects of the zinc/manganese (Zn/Mn) addition used to induce *ftsX* expression, we measured the growth of the parent and the FtsX merodiploid strain in the presence of these metals. To verify that the addition of 0.45 mM ZnCl_2_ and 0.045 mM MnSO_4_ (Zn/Mn) did not cause growth or morphological defects, cells were grown in the presence and absence of Zn/Mn and imaged at 3 h and 6 h into the growth curve (Fig. S7C-D). Wild-type cells (D39 ∆*cps rpsL1*) had no morphological or growth defects at these time points with or without the addition of Zn/Mn (Fig. S7C-E).

In contrast, the FtsX merodiploid strain (P_zn_-*ftsX* ∆*ftsX*) had significant morphological and growth defects at 3 or 6 h in the absence of Zn/Mn (Fig. S7C-E). Cessation of growth and aberrant cell morphology was observed in 90% of cells at 3 h and 95% of cells at 6 h growth (Fig. S7D). These cells are significantly shorter and rounder than wild-type cells (Fig. S7E), and a large variability in their volumes was observed (Fig. S7E), as previously found for a strain expressing *ftsX*^+^under a fucose-inducible promoter (21). If the strain was grown in the presence of Zn/Mn, FtsX was expressed and the strain has no growth or morphological defects (Fig. S7C-E). This indicates the defects observed are solely due to the absence of FtsX.

We next constructed three classes of amino acid substitution or insertion mutants (Table 4) in an effort to disrupt the FtsX_ECL1_-PcsB-CC interaction defined by NMR spectroscopy. These are designated Class I (single amino acid changes), Class II (multiple amino acid changes), and Class III (insertion of a (Gly_3_Ser)_2_ linker) mutants. Class I strains were made by introducing single amino acid substitutions in the merodiploid strain and measuring growth or morphology defects (Fig. S8, Table 4). Class I mutants targeted both the α2 helix and the loop region (residues 107–120, and 121–130, respectively) of the FtsX_ECL1_ helical lobe (Table 4, Fig. 3A, S8A). Single amino acid substitutions of FtsX (E109A, L115A, M119A, W123A, F126A, N131A, N131D) exhibited morphological defects without Zn/Mn (Fig. S8C-D, Table 4), but did not induce a measurable growth phenotype (Fig. S8B and Table 4). Expression of FtsX (L115A) resulted in cell shape defects (aspect ratio, length, width, and volume) (Fig. S8C-D), while expression of FtsX (M119A) only resulted in a change in cell volume (Fig. S8D). These differences were not due to mis-expression of FtsX, as western blotting indicates that all were expressed at or near wild-type levels (Fig. 4C). Two other single amino acid substitutions (E109Q and Q111A) did not strongly affect growth, morphology, or expression (Table 4, Fig. 4D).

**Figure 3:**
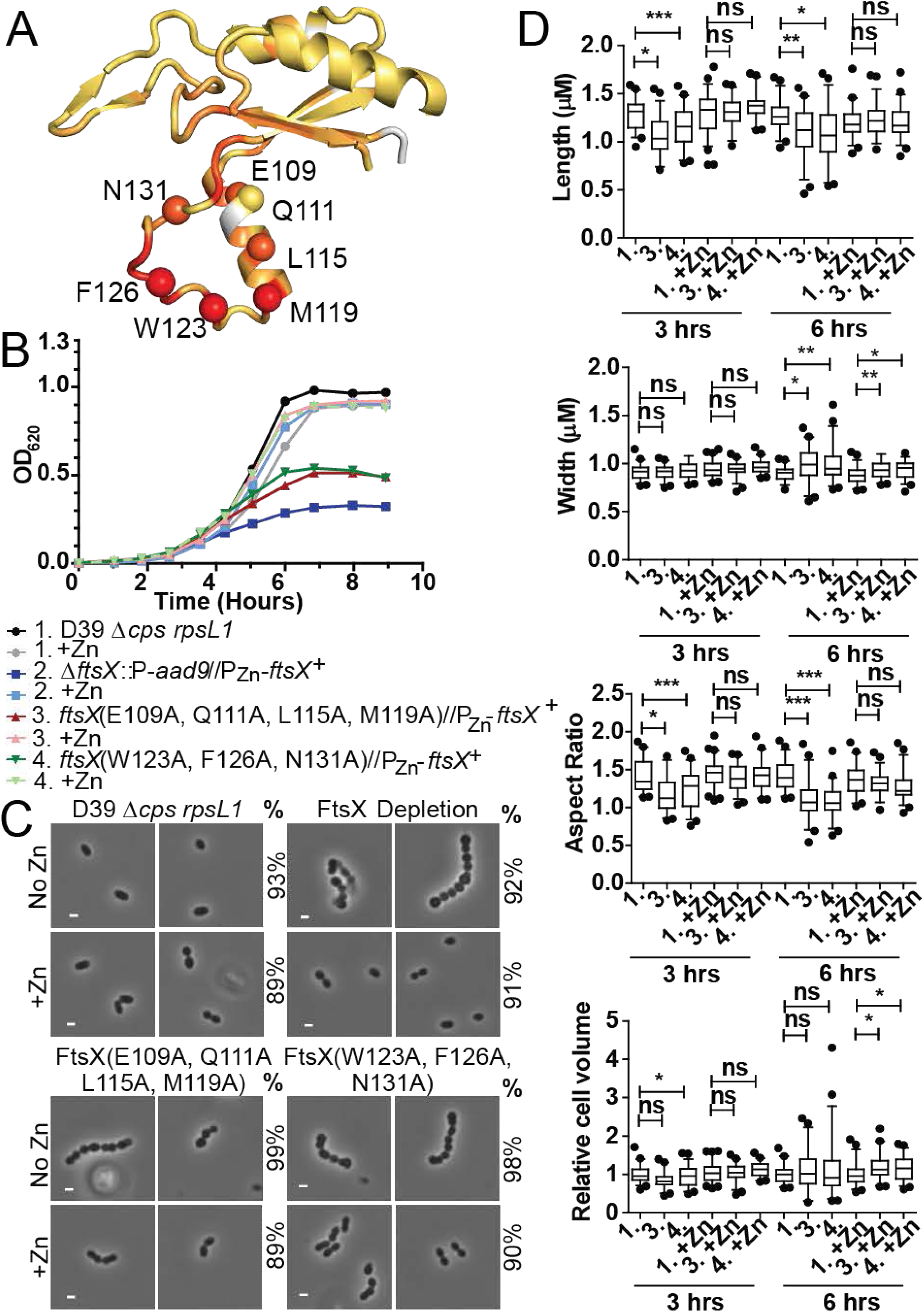
Multiple amino acid changes in the lower lobe of FtsX_ECL1_ cause morphological and growth defects. A) Amino acid changes made mapped to the structure of FtsX_ECL1_. The a carbon of each residue is shown as a colored sphere. The orange to red coloring on the FtsX_ECL1_ structure represents the peak height changes from ^1^H-^15^N HSQC spectra upon addition of 2 molar equivalents of PcsB-CC to FtsX_ECL1_ B) Representative growth curve of strains with amino acid changes in the lower lobe of FtsX_ECL1_, compared to the growth of a FtsX depletion strain. These strains were grown in the presence or absence of 0.45 mM ZnCl_2_ supplemented with 0.045 mM MnS04 (indicated as +Zn). Strains shown are as follows: black circle, D39 *rpsL1 Δcps* wild-type parent (1, IU1824); grey circle, IU1824 +Zn; dark blue square, D39 *rpsL1 Δcps ΔftsX.:P-aad9//bgaA::tet-*P_Zn_*-ftsX+* (2, IU12376); light blue square, IU12376 +Zn; red triangle, D39 *rpsL1 Δcps ftsX*(E109A, Q111A, L115A, M119A)*//bgaA*::*tet-*P_Zn_*-ftsX+* (3, IU12861); pink triangle, IU12861 +Zn; dark green inverted triangle, D39 *rpsL1 Δcps ftsX*(F126A, W123A, N131A)//*bgaA*::*tet*-P_zn_-*ftsX*+(4, IU12864); light green inverted triangle, IU12864 +Zn. This growth curve was repeated three times with similar results. C) Representative images of strains at 6 hours growth. The genotype of the strain shownis indicated above each panel. No Zn or +Zn indicates if Zn/Mn mixture was added. % indicates the percentage of cells in the population that are morphologically similar to the images shown. Greater than 50 cells per strain, condition, and experimental repeat were analyzed. These experiments were performed three times independently with similar results. Scale bar shown is equal to 1 pM. D) Length, width, aspect ratio, and relative cell volume of strains at 3 hours and 6 hours growth. Strains are indicated according to numbering in panel B. Greater than 50 cells were measured per strain and condition over two experimental replicates. For statistical analysis, a Kruskal-Wallis test (one-way AN OVA) with Dunn’s multiple comparison post-test was used to determine if length, width, aspect ratio, and relative cell volume were significantly differenl between strains and conditions. ns=non significant, * = p<0.05, ** = p<0.005, *** = p<0.0005

**TABLE 4.** Amino acid changes made *in vivo* to disrupt the FtsX_ECL1_-PcsB interaction.^a^

| Categories of Amino Acid Changes Made |  |  |  |
| --- | --- | --- | --- |
|  | Amino Acid Change | Defect in Shape and Growth | Location in FtsX<sub>ECL1</sub> |
| Class I | E109A | Shape Only | α2 helix |
|  | E109Q | No | α2 helix |
|  | Q111A | No<sup>b</sup> | α2 helix |
|  | L115A | Shape Only | α2 helix |
|  | M119A | Shape Only | α2 helix |
|  | W123A | Shape Only | Loop<sup>c</sup> |
|  | F126A | Shape Only | Loop<sup>c</sup> |
|  | N131A | Shape Only | Loop<sup>c</sup> |
|  | N131D | Shape Only | Loop<sup>c</sup> |
| Class II | E109A, N131A | Shape Only | α2 helix, Loop<sup>c</sup> |
|  | Q111A, L115C | Shape Only | α2 helix |
|  | L115A, M119A | Yes | α2 helix |
|  | E109A, Q111A, L115A, M119A | Yes | α2 helix |
|  | F126A, W123A, N131A | Yes | Loop<sup>c</sup> |
| Class III<br>(Gly<sub>3</sub>Ser)<sub>2</sub><sup>d</sup> | Residue 51 | Yes | post-TM1 |
|  | Residue 78 | Shape Only | β-hairpin |
|  | Residue 173 | N/A<sup>e</sup> | pre-TM2 |

<sup>a</sup>See Table S1 for corresponding strain number and strain construction for each amino acid change or insertion. <sup>b</sup>In the absence of zinc, this strain had no morphology defects. Morphology defects were observed at 3 h when the strain was expressing both the wild-type copy of FtsX and FtsX(Q111A). <sup>c</sup>Loop refers to the unstructured loop after $\alpha 2$ helix in the helical lobe of the FtsX<sub>ECL1</sub> structure. <sup>d</sup>These strains an insertion of the amino acid sequence GGGSGGGS after the indicated residue. <sup>e</sup>N/A: This strain did not express FtsX with (Gly<sub>3</sub>Ser)<sub>2</sub> insertion. See Fig. 4E for Western blot data.

In contrast to the somewhat modest physiological impact of Class I substitutions, selected Class II mutants (Table 4) exhibited severe morphological and growth defects (Fig. 3 and S9, Table 4). In strains targeting the α2 helix, FtsX(E109A, Q111A, L115A, M119A), or the coil, FtsX(F126A, W123A, N131A), ≥98% of cells had severe growth and morphology defects. The growth and morphology of these strains was similar to *ftsX* depleted cells (Fig. 3B-D). Cells expressing these *ftsX* alleles became significantly rounder and shorter (Fig. 3D), and growth was inhibited without zinc (Fig. 3B). Importantly, in the presence of zinc the cells were indistinguishable from wild-type at 3 h (Fig. 3D). At 6 h with zinc, these cells exhibited changes in width and volume, which could be due to overexpression of FtsX at this time point or expression of wild-type and mutant FtsX simultaneously (Fig. 3D). Western blotting confirmed that FtsX(E109A, Q111A, L115A, M119A) and FtsX(F126A, W123A, N131A) were expressed in the absence of zinc at 6 h post-depletion (Fig. 4A). The triple FtsX(F126A, W123A, N131A) mutant migrated slightly higher on an SDS-PAGE gel compared to wild-type FtsX in the absence of zinc, but was expressed (Fig. 4A). Taken together, these results reveal that both the α2 helix and loop in the helical lobe of FtsX_ECL1_ are important for FtsX function *in vivo* and confirm the functional importance of the physical interaction of FtsX and PcsB mapped by NMR.

**Figure 4:**
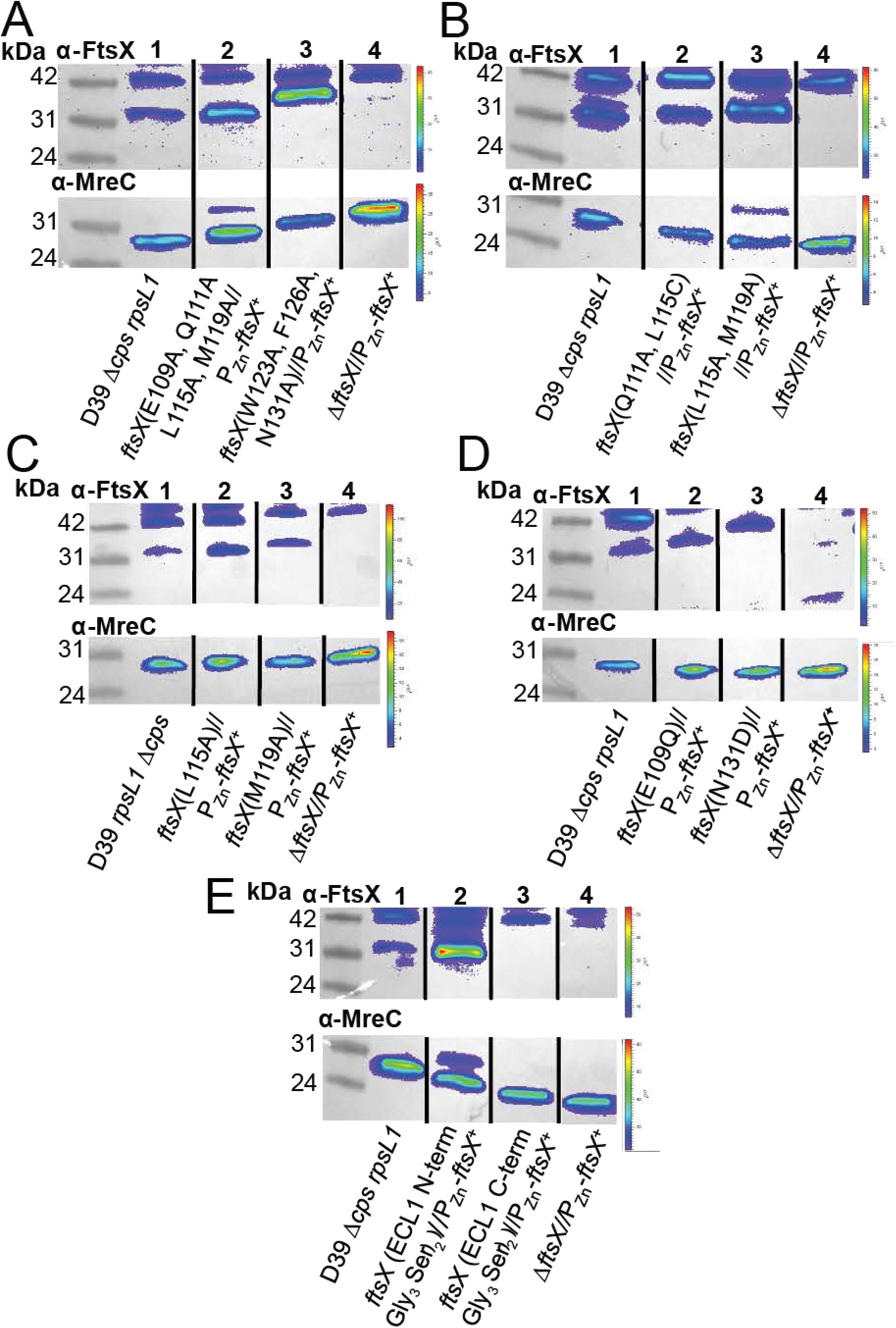
FtsX with amino acid changes are expressed at near wild-type levels. Representative blots of a-FtsX and α-MreC (Western blotting control for loading) are shown, with the genotype indicated under each lane. Expected molecular weight (MW) for FtsX is 34.2 kDa, expected MW for MreC is 29.7 kDa. Samples were grown without Zn and harvested at 6 hours growth. Westerns were imaged as described in *Supplemental Materials and Methods.* A) FtsX(E109A, Q111A, L115A, M119A) and FtsX(W123A, F126A, N131A) are expressed at or above wild-type levels without zinc. Lane 1, D39 *rpsL1 Δcps* (IU1824); lane 2, D39 *rpsL1 Δcps* ftsX(E109A, Q111A, L115A M119A)*//bgaA*::*tet*-P_zn_-*ftsX*+ (IU12861); lane 3, D39 *rpsL1 Δcps* ftsX(W123A, F126A, N131A)//*bgaA:.tet-*P_zn_*-ftsX+* (IU12864); lane 4, D39 *rpsL1 Δcps ΔftsX//bgaA:.tet-*P_zn_*-ftsX+* (IU13461). B) FtsX(Q111 A, L115C) and FtsX(L115A, M119A) are expressed at near wild-type levels without zinc. Lane 1, D39 *rpsL1 Δcps* (IU1824); lane 2, D39 *rpsL1* Acps ftsX(Q111A, L115C)//*bgaA*::*tet*-P_zn_-*ftsX*+ (IU13064); lane 3, D39 *rpsL1 Δcps ftsX*(L115A, M119A)//*bgaA*::*tet*-P_zn_-*ftsX*+ (IU13066); lane 4, D39 *rpsL1 Δcps ΔftsX//bgaA::tet-*P_zn_*-ftsX+.* C) FtsX (L115A) and FtsX (M119A) are expressed at near wild-type levels without zinc. Lane 1, D39 *rpsL1 Δcps* (IU1824); lane 2, D39 *rpsL1 Δcps ftsX*(L11A)*//bgaA::tet-*P_zn_*-ftsX+* (IU12521); lane 3, D39 *rpsL1 Δcps ftsX*(M119A)//*bgaA*::*tet*-P_zn_-*ftsX*+ (IU12637); lane 4, D39 *rpsL1 Δcps ΔftsX//bgaA::tet-*P_zn_*-ftsX+* (IU13461). D) FtsX (E109Q) and FtsX (N131D) are expressed at near wild-type levels without zinc. Lane 1, D39 *rpsL1 Δcps* (IU 1824); lane 2, D39 *rpsL1 Δcps ftsX*(E109Q)*//bgaA::tet-*P_zn_*-ftsX+* (IU13088); lane 3, D39 *rpsL1 Δcps ftsX*(*N*131D)*//bgaA::tet-*P_zn_*-ftsX+* (IU13089); lane 4, D39 *rpsL1 Δcps ΔftsX//bgaA::tet-*P_zn_*-ftsX+* (IU13461). E) FtsX with (Gly_4_Ser)_2_ after residue 51 is expressed, whereas FtsX with (Gly_4_Ser)_2_ after residue 173 is not expressed. These are referred to as FtsX N-term ECL1 (Gly_4_Ser)_2_ and FtsX C-terminal ECL1 (Gly_4_Ser)_2_, respectively. Lane 1, D39 *rpsL1 Δcps* (IU 1824); lane 2, D39 *rpsL1 Δcps ftsX N-term ECL1* (*Gly_4_Sery*)_2_*//bgaA::tet-*P_zn_*-ftsX+* (IU12629); lane 3, D39 *rpsL1 Δcps ftsX C-term ECL1* (*Gly_4_ Ser*)_2_*//bgaA.:tet-*P_zn_*-ftsX+* (IU12869); lane 4, D39 *rpsL1 Δcps ΔftsX/l bgaA:.tet-*P_zn_*-ftsX+* (IU 13461). These experiments were performed two to three times independently.

Some of the Class II mutants we characterized had just two amino acid changes in the α2 helix or the extended loop of the helical lobe (Table 4, Fig. S9). We observed that FtsX (L115A, M119A) exhibited a strong growth and morphology defect in the absence of zinc in 99% of cells at 6 h (Fig. S9B-C, Table 4), and these cells displayed a decrease in length, width, and volume relative to wild-type cells (Fig. S9D). This mutant was expressed in the absence of zinc at near wild-type levels (Fig. 4B). These data confirm that the tandem L115A, M119A substitution disrupts FtsX function, even though as individual single mutants they result only in slight morphological defects (Fig. S8B-D, Fig. S9B-D, Table 4). Another double mutant FtsX(Q111A, L115C) induced the formation of long chains and a “boxy” cell morphology (Fig. S9C, Table 4). This mutant resulted in shorter cells with a significantly different aspect ratio from wild-type (Fig. S9D), but this strain had no growth phenotype (Fig. S9B). In contrast, another double substitution Class II mutant, *e.g.,* FtsX(E109A, N131A) (Table 4) exhibited no strong morphology or growth defects.

Finally, Class III insertion mutants (Table 4) were constructed and used to evaluate if other regions of FtsX_ECL1_ were important for the FtsX_ECL1_-PcsB interaction or for FtsX function. We inserted a (Gly_3_Ser)_2_ flexible linker approximately where the FtsX_ECL1_ is predicted to enter (residue 51) or exit (residue 173) the membrane bilayer, or in the β-hairpin, which exhibits significant conformational disorder over a range of timescales (Fig. 1A). An insertion after residue 51 in FtsX_ECL1_ was detrimental to both growth and morphology (Table 4), and this insertion did not disrupt FtsX expression (Fig. 4E). The insertion after residue 173 in FtsX_ECL1_ also caused growth and morphology defects, but this FtsX allele was not expressed in cells (Table 4, Fig. 4E). The β-hairpin (Gly_3_Ser)_2_ insertion (Fig. 1B, Appendix) was introduced after amino acid 78 of FtsX_ECL1_, which corresponds to the tip of β-turn in the β-hairpin (Fig. 1B). This strain also exhibited no growth defect, but these cells were significantly smaller, but only at the 3 h time point (Appendix). We conclude that the β-hairpin does not play a major role in FtsX-PcsB interaction, consistent with the NMR mapping experiments.

### L115A/M119A FtsX_ECL1_ is stably folded and unable to bind PcsB-CC

We reasoned that if the defects observed in Class I and Class II mutants were due to the disruption of the FtsX_ECL1_-PcsB-CC interaction, this should impact the affinity of this interaction as measured by ITC. We first characterized the L115A/M119A mutant by ^1^H-^15^N NMR and CD spectroscopies to confirm its structural integrity. The CD spectrum resembled wild type, as did the ^1^H-^15^N HSQC spectrum, with clear chemical shift perturbations only among those resonances in the immediate vicinity of the double substitution (Fig. S10B). Both pieces of data suggest a local rather than global perturbation of α2-loop lobe in the FtsX_ECL1_ structure upon introduction of the double L115A/M119A substitution. As anticipated, titration of PcsB-CC (47–267) with L115A/M119A FtsX_ECL1_ reveals no detectable binding (no observable heat) (Fig. 2B, Table 3) compared to wild-type FtsX_ECL1_. These data confirm that the helical lobe of FtsX_ECL1_ interacts with PcsB-CC, and that this interaction is required for viability and proper cell shape.

In contrast to the L115A/M119A double mutant, severely functionally compromised representative triple mutant W123A, F126A, N131A FtsX_ECL1_ and quadruple mutant E109A, Q111A, L115A, M119A FtsX_ECL1_ exhibited more pronounced structural perturbations that nonetheless map only to the helical lobe. The triple mutant was indistinguishable from the L115A/M119A and wild-type FtsX_ECL1_ derivatives by CD spectroscopy while the quadruple mutant exhibited less molar ellipticity, or secondary structure (Fig. S10D). Inspection of their ^1^H-^15^N HSQC spectra reveal that although the core and β-hairpin domains essentially resemble wild-type, each exhibits considerable perturbation of resonances throughout the entire helical lobe (Fig. S10A, S10C). Since these mutants are functionally compromised, these structural findings strongly support the conclusion that the structural integrity of the lower lobe is essential for the physical interaction with PcsB, and the function of FtsX in pneumococcal cells.

## DISCUSSION

This study presents a comprehensive analysis of the solution and x-ray structures of the outward-facing large extracellular loop of FtsX (FtsX_ECL1_) from *S. pneumoniae* and defines a physical interaction site with the coiled-coil domain of peptidoglycan hydrolase PcsB (PcsB-CC). Our FtsX_ECL1_ structures reveal a globular fold that while similar to the large extracellular loop of FtsX from *M. tuberculosis* (16) is characterized by unique features. The upper β-hairpin distinguishes *S. pneumoniae* FtsX_ECL1_ from that of *M. tuberculosis*, and despite being characterized by significant conformational dynamics on a range of timescales, is not required for the interaction of *S. pneumoniae* FtsX_ECL1_ with PcsB. The function of this domain is not well-defined by our data, but could play a role in another process, *e.g.,* FtsX dimerization, interaction with the small loop ECL2, or an interaction with another domain of PcsB. On the other hand, the helical lobe of FtsX_ECL1_, common to both *S. pneumoniae* and *M. tuberculosis* FtsX structures, is vital for the interaction PcsB *in vitro*, and that this interface is functionally important *in vivo*. Increasing numbers of Ala substitutions tested here increasingly disrupt this interaction and ultimately cause dramatic growth and morphology defects, indicating that the helical lobe of FtsX_ECL1_ is essential for regulation of PcsB during cell division. Interestingly, this region of FtsX_ECL1_ corresponds with the region shown to be important for the interaction of *M. tuberculosis* FtsX_ECD_ with its PG hydrolase, RipC (16). This suggests that the helical lobe could be a conserved functional determinant for the interaction of FtsX with cognate hydrolases or adaptor proteins across many species of bacteria.

We propose that the helical lobe of FtsX_ECL1_ is important for the activation of cognate hydrolase activity either directly or indirectly through their adaptors (Fig. 5). The exact role of the second extracellular loop of FtsX (FtsX_ECL2_) is unknown, but may also regulate this process, as temperature-sensitive mutations in *pcsB* were found to be suppressed by mutations in the coding region for FtsX_ECL2_ (20). Previous work suggests that FtsEX forms a dimer (32), as dimerization of the FtsE ATPase domain is likely a necessary condition for ATP hydrolysis (36, 37) (Fig. 5A). After formation of the complex, ATP hydrolysis by FtsE results in a conformational change in FtsX, releasing PcsB from what we anticipate is an inhibited state (22) (Fig. 5B-C). This interaction is mediated by the helical lobe of FtsX_ECL1_, although the membrane, and possibly lipid binding by FtsX_ECL1_ and FtsX_ECL2_ itself also may play a role as well. We propose that the interaction of FtsX_ECL1_ with the PcsB coiled-coil domain communicates release of the PcsB CHAP domain from an inhibited state and thus is important for modulating PG hydrolysis by PcsB (Fig. 5B-C).

**Figure 5:**
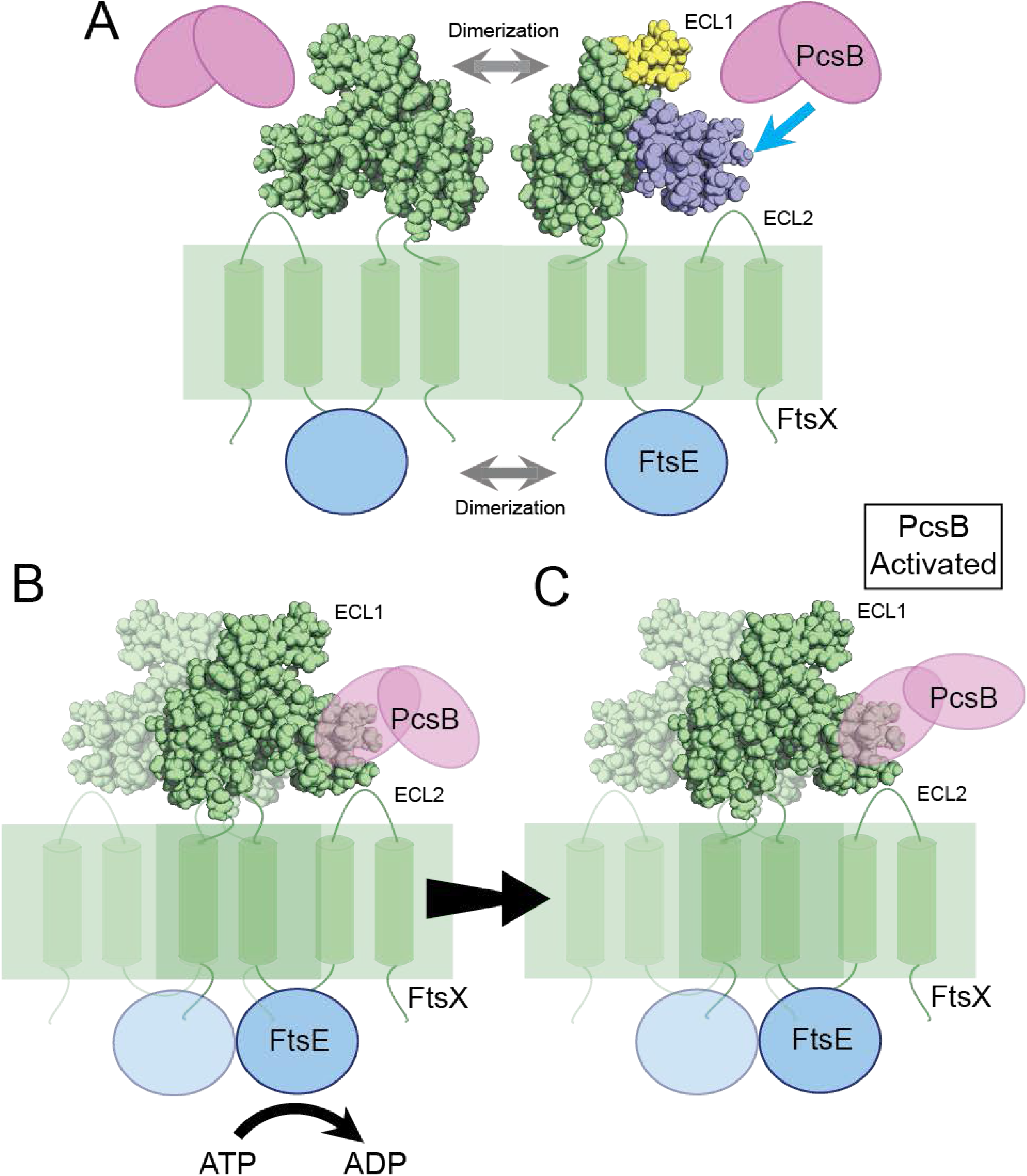
Model for the activation of PcsB by FtsX_ECL1_. A) FtsEX dimerizes to form the active complex. PcsB is secreted into the extracellular milieu. Attraction of PcsB to the area of active FtsX complexes might be mediated by its propensity to interact with membranes (Bajaj et. al, 2017). The ECL1 and ECL2 loops are indicated on FtsX. FtsXE_ECL1_ is shown in green, with the β-hairpin and α-helical lobe shaded in yellow and blue, respectively. B) After formation of the active complex, ATP hydrolysis by FtsE causes a conformational change in FtsX. PcsB interacts with FtsX_ECL1_ via its coiled coil domain, and this interaction causes activation of the peptidoglycan hydrolytic activity of PcsB. PcsB, along with other factors in the cell, allow for cell division to proceed normally. Functional FtsX, FtsE, and PcsB are all required for efficient cell division.

Recently, the structure of *A. actinomycetemcomitans* MacB was reported and suggested to be a structural paradigm for the ABC3 transporter superfamily that includes FtsX (38). MacB was proposed to function as a mechanical pump to drive enterotoxin transport through TolC in *E. coli* (38). Crow *et. al* found that MacB itself did not transport enterotoxin but drove TolC to transport it instead, due to the lack of a central cavity in the MacB structure (38). As such, they proposed that MacB as a model for other “mechanotransmitters” belonging to this same ABC3 transporter superfamily. While this proposed function of mechanotransmission may well characterize MacB and FtsX, the overall structure of FtsX_ECL1_ from *S. pneumoniae* is clearly distinct from the periplasmic domain of MacB, thus revealing that MacB does not readily provide a structural basis for understanding FtsX-dependent peptidoglycan hydrolases. Future work using reconstituted FtsEX and PcsB complexes in membranes will allow for understanding how this common mechanotransmission principle extends throughout the ABC3 superfamily.

## MATERIALS AND METHODS

### NMR Spectroscopy

Spectra of ^15^N-or ^15^N^13^C-labeled FtsX_ECL1_ were recorded at 298 K on Varian (Agilent) DDR 600 or 800 MHz spectrometers equipped with cryogenic probes in the METACyt Biomolecular NMR Laboratory at Indiana University Bloomington. NMR samples contained 50 mM potassium phosphate, pH 7.0, 50 mM NaCl and 10% v/v D2O, with 0.2 mM 4,4-dimethyl-4-silapentane-1-sulfonic acid (DSS) for chemical shift referencing. Typical concentrations of FtsX_ECL1_ for ^15^N HSQC spectra were 50 µM, and 400 µM for triple-resonance and dynamics experiments. ^1^*J*_HN_ splittings for residual dipolar couplings (RDCs) were measured using 2D IPAP [^15^N, ^1^H]-HSQC spectra (39), recorded on an isotropic sample and on a sample aligned with 20 mg/mL phage *Pf*1 (ASLA Biotech). ^1^*D*_HN_ was calculated from ^1^*D*_HN_ =^1^*J*_HN_ (anisotropic) – ^1^*J*_HN_ (isotropic). Aromatic sidechains were assigned using the HBCBCGCDHE and HBCBCGCDCEHE experiments (40). For experiments detecting PcsB-CC binding, ^15^N FtsX_ECL1_ was kept at 50 µM concentration, and ^1^H-^15^N HSQC spectra were recorded with the following concentrations of PcsB-CC (47–267): 0, 50 µM, 100 µM, 200 µM, and 400 µM. nmrPipe, Sparky, CARA (http://cara.nmr.ch), CCPNMR, and NMRbox (41–44) were used for data processing and analysis. Resonance assignments and dynamics data are available in the BMRB under accession code 30523. These NMR data were used to calculate and refine the solution structure of ECL1 (see *Supplemental Materials and Methods*) with the ensemble of 20 lowest energy structures (see Table 1 for structure statistics) deposited in the Protein Data Bank (accession code 6MK7).

### X-Ray Crystallography

First crystallization screenings were performed by high-throughput techniques in a NanoDrop robot and Innovadyne SD-2 microplates (Innovadyne Technologies Inc.), screening PACT Suite and JCSG Suite (Qiagen), JBScreen Classic 1–4 and 6 (Jena Bioscience) and Crystal Screen, Crystal Screen 2 and Index HT (Hampton Research). Positive conditions in which crystals grew were optimized by sitting-drop vapor-diffusion method at 291 K by mixing 1 µL of protein solution and 1 µL of precipitant solution, equilibrated against 150 µL of precipitant solution in the reservoir chamber. The best crystals were obtained in a crystallization condition containing 0.1 M sodium citrate pH=5.6, 0.2 M potassium-sodium tartrate and 2 M ammonium sulfate. These crystals were further optimized by cocrystallization in the presence of detergents Dodecyltrimethylammonium chloride (detergent 1, 46 µM), n-Undecyl-β-D-maltoside (detergent 2, 0.59 mM), and n-Decyl-β-D-maltoside (detergent 3, 1.8 mM). Crystals were cryo-protected in the precipitant solution supplemented with 25% (v/v) glycerol, prior to flash cooling at 100 K. Diffraction data were collected in beamline XALOC at the ALBA synchrotron (CELLS-ALBA, Spain), using a Pilatus 6M detector and a wavelength of 0.979257 Å. Crystals diffracted up to 2.0–2.3 Å resolution and belonged to the P 43 21 2 space group. The collected datasets were processed with XDS (45) and Aimless (46). Two FtsX_ECL1_ molecules were found in the asymmetric unit, yielding a Matthews coefficient of 2.49 Å^3^/Da (47) and a solvent content of 50.73%. Structure determination was performed by molecular replacement using the online server Morda (http://www.biomexsolutions.co.uk/morda/). Refinement and manual model building were performed with Phenix (48) and Coot (49), respectively. The data presented translational non-crystallographic symmetry that were treated with Phenix (48). The stereochemistry of the final model was checked by MolProbity (50). Data collection and processing statistics are shown in Table 2. The atomic coordinates of FtsX_ECL1_ determined by co-crystallization in the presence of detergents 1, 2 and 3 have been deposited in the Protein Data Bank with codes 6HE6, 6HEE and 6HFX, respectively.

### Bacterial strains, plasmids, and growth conditions

Bacterial strains and plasmids used in this study are listed in Table S1. *S. pneumoniae* strains were derived from IU1945, an unencapsulated derivative of serotype 2 *S. pneumoniae* strain D39 (51). Strains were grown on trypticase soy agar II with 5% (vol/vol) defibrinated sheep blood (TSAII-BA) plates or in Becton-Dickinson brain heart infusion (BHI) broth at 37°C in an atmosphere of 5% CO_2_. *E. coli* strains for protein expression were derived from BL21(DE3) (NEB, C2527H). *E. coli* strains were grown in Luria-Bertani (LB) broth or in M9 minimal media supplemented with ^15^NH_4_Cl at 37°C, with shaking at 150 rpm. When required, tetracycline (0.25–2.5 µg/mL), kanamycin (250 µg/mL), spectinomycin (150 µg/mL), streptomycin (250 µg/mL), ampicillin (100 µg/mL) and/or isopropyl β-D-1-thiogalactopyranoside (IPTG, 1 mM) were added to *S. pneumoniae* or *E. coli* culture media. *S. pneumoniae* strains requiring zinc for expression of essential genes were grown with 0.45 mM ZnCl_2_ and 0.045 MnSO_4_.

### Growth curves and phase-contrast microscopy of strains

For physiological and morphological analyses of strains, cells were inoculated from frozen glycerol stocks into BHI broth, serially diluted, and incubated for 10–12 hours statically at 37°C in 5% CO2 overnight. If zinc was required for growth of cultures, 0.45 mM ZnCl_2_ and 0.045 MnSO_4_ was added to overnight tubes. The next day, cultures from OD_620_ ≈ 0.05 to 0.4 were diluted into fresh BHI to OD_620_ ≈ 0.003 in 4 mL volumes, and two identical cultures for each strain were prepared, one with 0.45 mM ZnCl_2_/0.045 MnSO_4_ and one without. These cultures were grown under the same growth conditions as previously described in *Materials and Methods*. Growth was monitored turbidimetrically every 45 min to 1 hour with a Genesys 2 spectrophotometer (Thermo Scientific). For microscopic analyses, samples (1–2 μL) were taken at 3 hours and 6 hours and examined using a Nikon E-400 epifluorescence phase-contrast microscope with a 100X Nikon Plan Apo oil-immersion objective (numerical aperture, 1.40) connected to a CoolSNAP HQ2 charged-coupled device (CCD) camera (Photometrics). Images were processed using NIS-Elements AR software (Nikon), and measurements and calculation of cell width, length, volume, and aspect ratio were performed as described previously (52, 53). Statistical significance was determined using GraphPad Prism (GraphPad Software, Inc) by comparing values for cell width, length, volume, and aspect ratio measured for at least 50 cells over two experimental replicates. To determine if values were significantly different between strains and conditions, a Kruskal-Wallis (one-way ANOVA) with Dunn’s multiple comparison post-test was used.

**For additional materials and methods, please see Supplemental Materials and Methods.**

## ACKNOWLEDGEMENTS

We thank the members Winkler and Giedroc labs for their helpful discussion and insight, specifically Dr. Tiffany Tsui, Dr. Julia Martin, Melissa Lamanna and Dr. Hui Peng. We acknowledge the Indiana University Bloomington Department of Chemistry Mass Spectrometry Facility, specifically Dr. Jonathan Karty and Angela Hansen for their help with training and setting up mass spectrometry experiments and the MetaCyt Biomolecular NMR Laboratory. We also thank Dr. Daiana Capdevila for her help in acquiring and analyzing the isothermal titration calorimetry data, the Indiana University Biological Mass Spectrometry Facility and Dr. Giovanni Gonzales-Gutierrez in the Physical Biochemistry Instrumentation Facility at Indiana University Bloomington. We thank the staff from ALBA synchrotron facility for help during crystallographic data collection. This work was supported by NIH grants R01GM114315 and R01GM127715 to M.E.W., and NIH grant R35GM118157 to D.P.G. and by predoctoral Quantitative and Chemical Biology (QCB) NIH institutional training grant T32 GM109825 (to B.E.R.). The work in Spain was supported by grant from the Spanish Ministry of Science, Innovation and Universities BFU2017-90030-P to J.A.H.

## REFERENCES

1. Henriques-Normark B, Normark S. 2010. Commensal pathogens, with a focus on Streptococcus pneumoniae, and interactions with the human host. Exp Cell Res 316: 1408–14.

2. Kadioglu A, Weiser JN, Paton JC, Andrew PW. 2008. The role of Streptococcus pneumoniae virulence factors in host respiratory colonization and disease. Nat Rev Microbiol 6: 288–301.

3. Chao Y, Marks LR, Pettigrew MM, Hakansson AP. 2014. Streptococcus pneumoniae biofilm formation and dispersion during colonization and disease. Front Cell Infect Microbiol 4: 194.

4. Feldman C, Anderson R. 2014. Recent advances in our understanding of Streptococcus pneumoniae infection. F1000Prime Rep 6: 82.

5. CDC. 2013. Antibiotic resistance threats in the United States, 2013. http://www.cdc.gov/drugresistance/threat-report-2013/,

6. Garcia-Bustos J, Tomasz A. 1990. A biological price of antibiotic resistance: major changes in the peptidoglycan structure of penicillin-resistant pneumococci. Proc Natl Acad Sci U S A 87: 5415–9.

7. Massidda O, Novakova L, Vollmer W. 2013. From models to pathogens: how much have we learned about Streptococcus pneumoniae cell division? Environ Microbiol 15: 3133–57.

8. Sham LT, Tsui HC, Land AD, Barendt SM, Winkler ME. 2012. Recent advances in pneumococcal peptidoglycan biosynthesis suggest new vaccine and antimicrobial targets. Curr Opin Microbiol 15: 194–203.

9. Egan AJ, Cleverley RM, Peters K, Lewis RJ, Vollmer W. 2017. Regulation of bacterial cell wall growth. FEBS J 284: 851–867.

10. Vollmer W, Blanot D, de Pedro MA. 2008. Peptidoglycan structure and architecture. FEMS Microbiol Rev 32: 149–67.

11. Rajagopal M, Walker S. 2017. Envelope Structures of Gram-Positive Bacteria. Curr Top Microbiol Immunol 404: 1–44.

12. Rojas ER, Huang KC. 2018. Regulation of microbial growth by turgor pressure. Curr Opin Microbiol 42: 62–70.

13. Holtje JV. 1998. Growth of the stress-bearing and shape-maintaining murein sacculus of Escherichia coli. Microbiol Mol Biol Rev 62: 181–203.

14. Holtje JV, Heidrich C. 2001. Enzymology of elongation and constriction of the murein sacculus of Escherichia coli. Biochimie 83: 103–8.

15. Alcorlo M, Martinez-Caballero S, Molina R, Hermoso JA. 2017. Carbohydrate recognition and lysis by bacterial peptidoglycan hydrolases. Curr Opin Struct Biol 44: 87–100.

16. Mavrici D, Marakalala MJ, Holton JM, Prigozhin DM, Gee CL, Zhang YJ, Rubin EJ, Alber T. 2014. Mycobacterium tuberculosis FtsX extracellular domain activates the peptidoglycan hydrolase, RipC. Proc Natl Acad Sci U S A doi:10.1073/pnas.1321812111.

17. Margulieux KR, Liebov BK, Tirumala V, Singh A, Bushweller JH, Nakamoto RK, Hughes MA. 2017. Bacillus anthracis Peptidoglycan Integrity Is Disrupted by the Chemokine CXCL10 through the FtsE/X Complex. Front Microbiol 8: 740.

18. Meier EL, Daitch AK, Yao Q, Bhargava A, Jensen GJ, Goley ED. 2017. FtsEX-mediated regulation of the final stages of cell division reveals morphogenetic plasticity in Caulobacter crescentus. PLoS Genet 13:e1006999.

19. Zielinska A, Billini M, Moll A, Kremer K, Briegel A, Izquierdo Martinez A, Jensen GJ, Thanbichler M. 2017. LytM factors affect the recruitment of autolysins to the cell division site in Caulobacter crescentus. Mol Microbiol 106: 419–438.

20. Sham LT, Jensen KR, Bruce KE, Winkler ME. 2013. Involvement of FtsE ATPase and FtsX extracellular loops 1 and 2 in FtsEX-PcsB complex function in cell division of Streptococcus pneumoniae D39. MBio 4.

21. Sham LT, Barendt SM, Kopecky KE, Winkler ME. 2011. Essential PcsB putative peptidoglycan hydrolase interacts with the essential FtsXSpn cell division protein in Streptococcus pneumoniae D39. Proc Natl Acad Sci U S A 108:E1061–9.

22. Bartual SG, Straume D, Stamsas GA, Munoz IG, Alfonso C, Martinez-Ripoll M, Havarstein LS, Hermoso JA. 2014. Structural basis of PcsB-mediated cell separation in Streptococcus pneumoniae. Nat Commun 5: 3842.

23. Meisner J, Montero Llopis P, Sham LT, Garner E, Bernhardt TG, Rudner DZ. 2013. FtsEX is required for CwlO peptidoglycan hydrolase activity during cell wall elongation in Bacillus subtilis. Mol Microbiol 89: 1069–83.

24. Yang DC, Peters NT, Parzych KR, Uehara T, Markovski M, Bernhardt TG. 2011. An ATP-binding cassette transporter-like complex governs cell-wall hydrolysis at the bacterial cytokinetic ring. Proc Natl Acad Sci U S A 108:E1052–60.

25. Yang DC, Tan K, Joachimiak A, Bernhardt TG. 2012. A conformational switch controls cell wall-remodelling enzymes required for bacterial cell division. Mol Microbiol 85: 768–81.

26. Du S, Pichoff S, Lutkenhaus J. 2016. FtsEX acts on FtsA to regulate divisome assembly and activity. Proc Natl Acad Sci U S A 113:E5052–61.

27. Schmidt KL, Peterson ND, Kustusch RJ, Wissel MC, Graham B, Phillips GJ, Weiss DS. 2004. A predicted ABC transporter, FtsEX, is needed for cell division in Escherichia coli. J Bacteriol 186: 785–93.

28. Reddy M. 2007. Role of FtsEX in cell division of Escherichia coli: viability of ftsEX mutants is dependent on functional SufI or high osmotic strength. J Bacteriol 189: 98–108.

29. Fu Y, Bruce KE, Rued B, Winkler ME, Giedroc DP. 2016. 1H, 13C, 15N resonance assignments of the extracellular loop 1 domain (ECL1) of Streptococcus pneumoniae D39 FtsX, an essential cell division protein. Biomol NMR Assign 10: 89–92.

30. Zweckstetter M. 2008. NMR: prediction of molecular alignment from structure using the PALES software. Nat Protoc 3: 679–90.

31. Ishima R. 2014. CPMG relaxation dispersion. Methods Mol Biol 1084: 29–49.

32. Bajaj R, Bruce KE, Davidson AL, Rued BE, Stauffacher CV, Winkler ME. 2016. Biochemical characterization of essential cell division proteins FtsX and FtsE that mediate peptidoglycan hydrolysis by PcsB in Streptococcus pneumoniae. Microbiologyopen 5: 738–752.

33. Sung CK, Li H, Claverys JP, Morrison DA. 2001. An rpsL cassette, janus, for gene replacement through negative selection in Streptococcus pneumoniae. Appl Environ Microbiol 67: 5190–6.

34. Jacobsen FE, Kazmierczak KM, Lisher JP, Winkler ME, Giedroc DP. 2011. Interplay between manganese and zinc homeostasis in the human pathogen Streptococcus pneumoniae. Metallomics 3: 38–41.

35. Martin JE, Lisher JP, Winkler ME, Giedroc DP. 2017. Perturbation of manganese metabolism disrupts cell division in Streptococcus pneumoniae. Mol Microbiol 104: 334–348.

36. Locher KP. 2016. Mechanistic diversity in ATP-binding cassette (ABC) transporters. Nat Struct Mol Biol 23: 487–93.

37. Rees DC, Johnson E, Lewinson O. 2009. ABC transporters: the power to change. Nat Rev Mol Cell Biol 10: 218–27.

38. Crow A, Greene NP, Kaplan E, Koronakis V. 2017. Structure and mechanotransmission mechanism of the MacB ABC transporter superfamily. Proc Natl Acad Sci U S A 114: 12572–12577.

39. Ottiger M, Delaglio F, Bax A. 1998. Measurement of J and dipolar couplings from simplified two-dimensional NMR spectra. J Magn Reson 131: 373–8.

40. Yamazaki T, Yoshida M, Nagayama K. 1993. Complete assignments of magnetic resonances of ribonuclease H from Escherichia coli by double-and triple-resonance 2D and 3D NMR spectroscopies. Biochemistry 32: 5656–69.

41. Lee W, Tonelli M, Markley JL. 2015. NMRFAM-SPARKY: enhanced software for biomolecular NMR spectroscopy. Bioinformatics 31: 1325–7.

42. Vranken WF, Boucher W, Stevens TJ, Fogh RH, Pajon A, Llinas M, Ulrich EL, Markley JL, Ionides J, Laue ED. 2005. The CCPN data model for NMR spectroscopy: development of a software pipeline. Proteins 59: 687–96.

43. Maciejewski MW, Schuyler AD, Gryk MR, Moraru, II, Romero PR, Ulrich EL, Eghbalnia HR, Livny M, Delaglio F, Hoch JC. 2017. NMRbox: A Resource for Biomolecular NMR Computation. Biophys J 112: 1529–1534.

44. Delaglio F, Grzesiek S, Vuister GW, Zhu G, Pfeifer J, Bax A. 1995. NMRPipe: a multidimensional spectral processing system based on UNIX pipes. J Biomol NMR 6: 277–93.

45. Kabsch W. 2010. Xds. Acta Crystallogr D Biol Crystallogr 66: 125–32.

46. Evans PR, Murshudov GN. 2013. How good are my data and what is the resolution? Acta Crystallogr D Biol Crystallogr 69: 1204–14.

47. Matthews BW. 1968. Some crystal forms of bovine chymotrypsinogen B and chymotrypsinogen A. J Mol Biol 33: 499–501.

48. Adams PD, Afonine PV, Bunkoczi G, Chen VB, Davis IW, Echols N, Headd JJ, Hung LW, Kapral GJ, Grosse-Kunstleve RW, McCoy AJ, Moriarty NW, Oeffner R, Read RJ, Richardson DC, Richardson JS, Terwilliger TC, Zwart PH. 2010. PHENIX: a comprehensive Python-based system for macromolecular structure solution. Acta Crystallogr D Biol Crystallogr 66: 213–21.

49. Emsley P, Lohkamp B, Scott WG, Cowtan K. 2010. Features and development of Coot. Acta Crystallogr D Biol Crystallogr 66: 486–501.

50. Chen VB, Arendall WB, 3rd, Headd JJ, Keedy DA, Immormino RM, Kapral GJ, Murray LW, Richardson JS, Richardson DC. 2010. MolProbity: all-atom structure validation for macromolecular crystallography. Acta Crystallogr D Biol Crystallogr 66: 12–21.

51. Lanie JA, Ng WL, Kazmierczak KM, Andrzejewski TM, Davidsen TM, Wayne KJ, Tettelin H, Glass JI, Winkler ME. 2007. Genome sequence of Avery’s virulent serotype 2 strain D39 of Streptococcus pneumoniae and comparison with that of unencapsulated laboratory strain R6. J Bacteriol 189: 38–51.

52. Land AD, Tsui HC, Kocaoglu O, Vella SA, Shaw SL, Keen SK, Sham LT, Carlson EE, Winkler ME. 2013. Requirement of essential Pbp2x and GpsB for septal ring closure in Streptococcus pneumoniae D39. Mol Microbiol 90: 939–55.

53. Tsui HC, Zheng JJ, Magallon AN, Ryan JD, Yunck R, Rued BE, Bernhardt TG, Winkler ME. 2016. Suppression of a deletion mutation in the gene encoding essential PBP2b reveals a new lytic transglycosylase involved in peripheral peptidoglycan synthesis in Streptococcus pneumoniae D39. Mol Microbiol 100: 1039–65.

